# Dried-bakery waste as a substrate for *n*-caproate and *n*-caprylate production *via* chain elongation: bakeroate

**DOI:** 10.64898/2025.12.20.695658

**Authors:** Jean Népomuscène Ntihuga, Joseph G. Usack, Raphael Mayer, Ahmed A. Kinbokun, Hatice Yesil, Mei Zhou, Largus T. Angenent

## Abstract

Bakery waste is a promising feedstock for the circular economy; however, its use for producing medium-chain carboxylates (MCCs) *via* microbial chain elongation remains unexplored. This study investigated pretreatment, pertraction, and chain elongation strategies to convert bakery waste into *n*-caproate and *n*-caprylate. Bakery waste was mechanically and enzymatically processed, and different inocula were tested to optimize the conversion of the resulting glucose-rich solution into lactate and ethanol (intermediates). These intermediates were fed into continuous chain elongation systems, which operated for over 386 days at 37°C and pH 5.5. Results showed that 30–35% of bakery waste carbon was converted into *n*-caproate and 10–15% into *n*-caprylate. Enzymatic starch hydrolysis proved essential, and lactate was a superior intermediate for chain elongation compared to ethanol. A maximum volumetric MCC production rate of 131 mM C L^-1^ d^-1^ (0.1 g L^-1^ h^-1^) was achieved. This integrated approach, named “bakeraote,” demonstrates an efficient pathway for valorizing bakery waste.

## 1. Introduction

The growing global human population has led to an increased demand for food. The rise in food production and wasteful consumer behaviour has also led to more food-related waste, especially from perishable and semi-perishable products. The imbalance between food production locations and waste generation has become an essential obstacle to food sustainability, resource efficiency, and preservation. A Food and Agriculture Organization (FAO) study revealed that approximately 1.3 billion tons of food waste are produced annually worldwide, equating to the release of 4.4 gigatons of CO_2_-eq into the environment and a financial loss of nearly $990 billion (Kim, B. C. *et al*., 2021). In many countries, short shelf-life and staple food products, such as bread, contribute considerably to food waste (Kumar *et al*., 2023). For instance, Europe alone produced 3.5 million tons of bakery waste in 2015, with 1.7 million tons originating from Germany (Borrull, 2015).

The optimal solution to reduce bakery waste includes addressing its sources and introducing best waste management practice policies (Brancoli *et al*., 2020). However, predicting consumer attitudes within the bakery-waste supply chain poses a challenge. Therefore, bakery waste will remain a problem. Fortunately, bakery waste represents an essential source of carbohydrates and substrates for waste valorization platforms that aim to produce valuable chemicals or bioenergy. Previous studies have shown that including bakery waste into bio-based waste management systems can result in the production of biochemicals *(e.g.,* 5-hydroxymethyl-2-furfurylamine), bioenergy (*e.g.,* ethanol and methane), and animal feed through microbial processes, which revalues this waste and reduces its environmental impact (Jung *et al*., 2022; Narisetty *et al*., 2022; Gao *et al*., 2023; Swetha *et al*., 2023). Despite technological advancements, many bio-based management systems rely mainly on governmental subsidies due to the product’s limited commercial value (*e.g.,* methane, ethanol). Consequently, there is a pressing need for a biotechnological platform that transforms bakery waste into more valuable chemicals such as *n-*caproate (4–6 $/kg) and *n-*caprylate (6–8 $/kg) (Debergh *et al*., 2022).

Microbial chain elongation is a carbon recovery and production platform that converts complex organic waste into medium-chain carboxylates (MCCs) (*i.e., n*-caproate, *n*-caprylate) as valuable chemicals. This process involves converting short-chain carboxylates (SCCs) and alcohols into MCCs through anaerobic fermentation with cooperative interactions between microbial populations in a food web. Demonstrated examples have highlighted the robustness of chain elongation in revalorizing organic residues, including acid whey (Xu *et al*., 2019), wastewater (Wu *et al*., 2020), food waste (Huo *et al*., 2024), municipal solid waste (Grootscholten *et al*., 2014), and gas substrates (Quintela *et al*., 2024). During fermentation, typical metabolic processes break down organic polymers into oligomers, which can be further fermented into chain elongation intermediates, namely lactate and ethanol. These intermediates are subsequently converted into MCCs through reverse ß-oxidation (Angenent *et al*., 2016). Bakery waste stands out as a highly promising substrate for microbial chain elongation due to its carbohydrate content, which can be converted into lactate and ethanol, and its proteins and amino acids, which can be taken as nitrogen sources for microbial growth. Moreover, using bakery waste for microbial chain elongation offers several advantages, yielding more valuable chemicals (*e.g*., *n*-caproate) than traditional management systems, which produce methane *via* anaerobic digestion (Battista *et al*., 2024).

Despite the high biotechnological potential of bakery waste as a substrate for microbial chain elongation, there have been no studies with this focus. Previous reports on chain elongation have focused on using other food wastes such as fruits, vegetables, and kitchen waste (Nzeteu *et al*., 2018; Contreras-Dávila *et al*., 2020; Kim, H. *et al*., 2022; Wang *et al*., 2022). Therefore, exploring bakery waste as a substrate for microbial chain elongation presents an opportunity to establish a sustainable and efficient platform.

The bakery-waste starch must be fully hydrolyzed into simple sugars before being used as a microbial chain elongation substrate, because most chain elongating bacteria cannot degrade macromolecules (Li *et al*., 2024). In addition, when the bakery waste starch is incompletely hydrolyzed, high levels of dextrin are formed, which result in a sticky mass (Every *et al*., 1996). The stickiness of bakery waste may also be due to damaged starch (Teobaldi *et al*., 2024). Therefore, careful pretreatment is required to avoid building up the sticky mass on the bioreactor walls and in the tubing due to the presence of the products, as mentioned above.

This study explored the potential of using bakery waste in a chain-elongation-production platform as a strategy for carbon recovery and waste management. The main areas of focus included: **(a)** evaluating and optimizing the conversion of bakery waste into chain-elongation intermediates; **(b)** evaluating the feasibility of using bakery-waste mash of different solid content in hollow-fiber membrane extraction applications; and **(c)** producing *n*-caproate and *n*-caprylate. The results of this study are intended to support the integration of organic wastes, particularly those with high carbohydrate content, into a biorefinery concept to produce higher-value products compared to typical waste management strategies.

## 2. Materials and Methods

We summarized the experimental methods in the main text. The electronic supplementary information (ESI) contains additional information on experimental parameters, methods, and results. We divided the conversion of bakery waste into the overall process flow into three steps: **(1)** the bakery waste pretreatment, which included bakery waste collection, transport, drying, size reduction, and hydrolysis; **(2)** the conversion of bakery waste into lactate/ethanol for subsequent chain elongation, which included optimizing lactate/ethanol production; and **(3)** the chain elongation process (**Fig. S1,** ESI †). Equations for calculating parameters are also provided in the ESI † (**Eq. S1-S8** in **Bioreactor equations**).

### 2.1. Pretreatment

#### 2.1.1. Bakery-waste pretreatment

We collected bakery waste with different chemical compositions (**Table S1,** ESI †) from Bäckerhaus Veit (Tübingen, Germany), which we first mechanically and then enzymatically processed. The mechanical processing included drying in ambient air and grinding with a bread grinder (MAC 100, MAC.PAN snc, Thiene, Italy). We used three grinding options in the bakery-waste grinder to achieve different particle size categories of bakery waste. Then, we measured the particle-size distribution of each category using the Mastersizer 2000 (Malvern Instruments Ltd, Worcestershire WR14 1XZ, United Kingdom). As a result, we characterized the ground dried-bakery waste into three categories: **(a)** small-particle dried bakery waste (particle size distribution ≤ 362 μm); **(b)** medium-particle dried bakery waste (particle size distribution between 360 and 680 μm); and **(c)** large-particle dried bakery waste (particle size distribution ≥ 680 μm). The enzymatic processing included liquefaction and saccharification in the Braumeister 20 L (Speidel Tank-und Behälterbau GmbH, Ofterdingen, Germany) using Amylase Thermo and Glucoamylase AN (*Aspergillus niger)* to convert the starch from the bakery-waste mash (8% w/v) into simple sugars. We performed gelatinization, liquefaction, and saccharification according to the enzyme manufacturer’s recommendations (ASA Spezialenzyme GmbH, Wolfenbüttel, Germany), resulting in small-particle mash, medium-particle mash, and large-particle mash.

#### 2.1.2. Lactate and ethanol production and optimization

We first optimized lactate and ethanol production considering the following: **(a)** mash pretreatment (small-particle mash, medium-particle mash, and large-particle mash); **(b)** temperature (25^°^C, 37^°^C, and 50^°^C); and **(c)** inoculum type. For the inoculum, we utilized three types: **(1)** a lactate microbiome, which is an inoculum containing *Lactobacillus* sp. with > 98% relative abundance (Schuetterle *et al*., 2024); **(2)** SiloSolve MC (microbial control), which is a commercial product and a mixture of *Lactobacillus plantarum* (CH 6072), *Enterococcus faecium* (DSM 22502), and *Lactococcus lactis* (SR3.54); and **(3)** *Bacillus coagulans* (DSM1). The bacterial culture *B. coagulans* DSM1 strain was procured from the German Collection of Microorganisms and Cell Cultures GmbH (DSMZ, Deutsche Sammlung von Mikroorganismen und Zellkulturen GmbH, Braunschweig, Germany) (**Fig. S1**, ESI †). We directly inoculated the first two inocula mentioned into a continuously-fed 1-L bioreactor (Klask *et al*., 2020) and pre-cultured the third inoculum on tryptone soya medium (**Tryptone soya medium**, ESI †) at a pH of 7.0 and a temperature of 50^°^C. For ethanol production, we used the technical yeast Kornbrand, referred to here as yeast (C. Schliessmann Kellerei-Chemie GmbH & Co. KG, Schwäbisch Hall, Germany). After optimizing lactate and ethanol production, we prepared 25 L of the small-particle fermented mash in batches and then mixed them to get a total volume of 1000 L for lactate and ethanol product (**Lactate and ethanol preparations**, ESI †). We stored the small-particle fermented mashes at-20^°^C until usage.

### 2.2 Chain elongation process

#### 2.2.1. Feasibility of using bakery-waste mash in liquid-liquid extraction in the chain elongation process

The extraction system is described in detail in the ESI † (**Description of extraction system**). Before operating the chain elongation system, which includes in-line extraction, we tested the feasibility of mash in membrane-based liquid-liquid extraction (*i.e*., pertraction) for the chain elongation process to ascertain its compatibility with this extraction system. We constructed a pertraction system that mimics the experimental setup of the chain elongation system (**Fig. S2,** ESI †). We first established a cake during a 12-h fouling experiment for the three different mashes by pushing each mash through the hollow-fiber membrane with a pump (raffinate rate was not constant) at a pressure of 0.6-0.9 bar. Subsequently, we performed a 12-h extraction experiment by connecting the fouled hollow-fiber membranes to the synthetic fermentation broth and the solvent in the same manner as the conventional pertraction system, which uses diffusion of undissociated carboxylic acids rather than pushing a liquid through the membrane. We circulated each of the three mashes, which were enriched with 10 g L^-1^ of *n*-caproate, through the shell side and the solvent through the lumen of the fiber at a flow rate of 2.2 L h^-1^ and 4.3 L h^-1^, respectively. We maintained a pressure of 0.5 bar on the shell side and 0.2 bar on the lumen side at a temperature of 37^°^C. We measured the extracted *n*-caproate in the solvent by weighing the solvent at the end of each batch (12 h), assuming that only *n*-caproate diffused. However, we needed to correct for the solvent that remained in the lumen by measuring the solvent reference weight. To measure the solvent reference weight, we first weighed the solvent before adding it to the lumen. Then, we circulated water on the shell side and the solvent for 2, 4, 6, 8, 10, and 12 h before weighing the solvent (six times) (**Batch extraction calculations,** ESI†). We successfully verified our approach by connecting the extraction system to a stripping system and measuring the *n*-caproate concentration in the stripping solutions (**Fig. S2**, ESI†).

#### 2.2.2. Chain elongation bioreactor system

We added yeast extract (1.25 g L^-1^) and trace elements to the small-particle fermented mash before feeding to the bioreactors, but without vitamins and minerals (**Bioreactor medium preparation**, ESI†) (Rajagopalan *et al*., 2002; Vasudevan *et al*., 2014; Kucek *et al*., 2016; Spirito *et al*., 2018). In addition, we added hydrochloric acid or sodium hydroxide to adjust the pH between 5.0 and 5.4. Depending on the operating period, we added tap water to dilute the small-particle fermented mash, and referred to the solutions as lactate medium or ethanol medium. We set up two independently, identically, and continuously fed upflow bioreactors with fermentation-broth recirculation through both a direct route and through a pertraction system (**Fig. 1**). One bioreactor was fed with lactate medium and the other bioreactor with ethanol medium. We used a peristaltic pump (Model 07528-20, Cole Parmer) to continuously supply fresh medium at room temperature from a 5-L medium tank to each bioreactor. This medium tank was flushed with nitrogen gas. We used peristaltic pumps (Model 7528-30, Cole-Parmer, Vernon Hills, IL, USA) to recirculate the fermentation broth through a pertraction system (**Description of extraction system,** ESI †). Each bioreactor was connected to an automated thermostat (CC 104A, Huber, Raleigh, NC, USA), which controlled the fermentation-broth temperature at 37^°^C for lactate and 30^°^C for ethanol (**Table S2,** ESI †). The extraction systems for each bioreactor were controlled at the same temperature as the fermentation broth. We placed the pH electrode (ProcessLine 80-325pH, SI Analytics, Weilheim, Germany) through the lid of the bioreactor and connected it to a pH controller (Bluelab pH controller, Bluelab Corporation Limited, Tauranga, New Zealand). The controller maintained the pH at 5.5 by adding hydrochloric acid (3 M).

**Figure 1.**
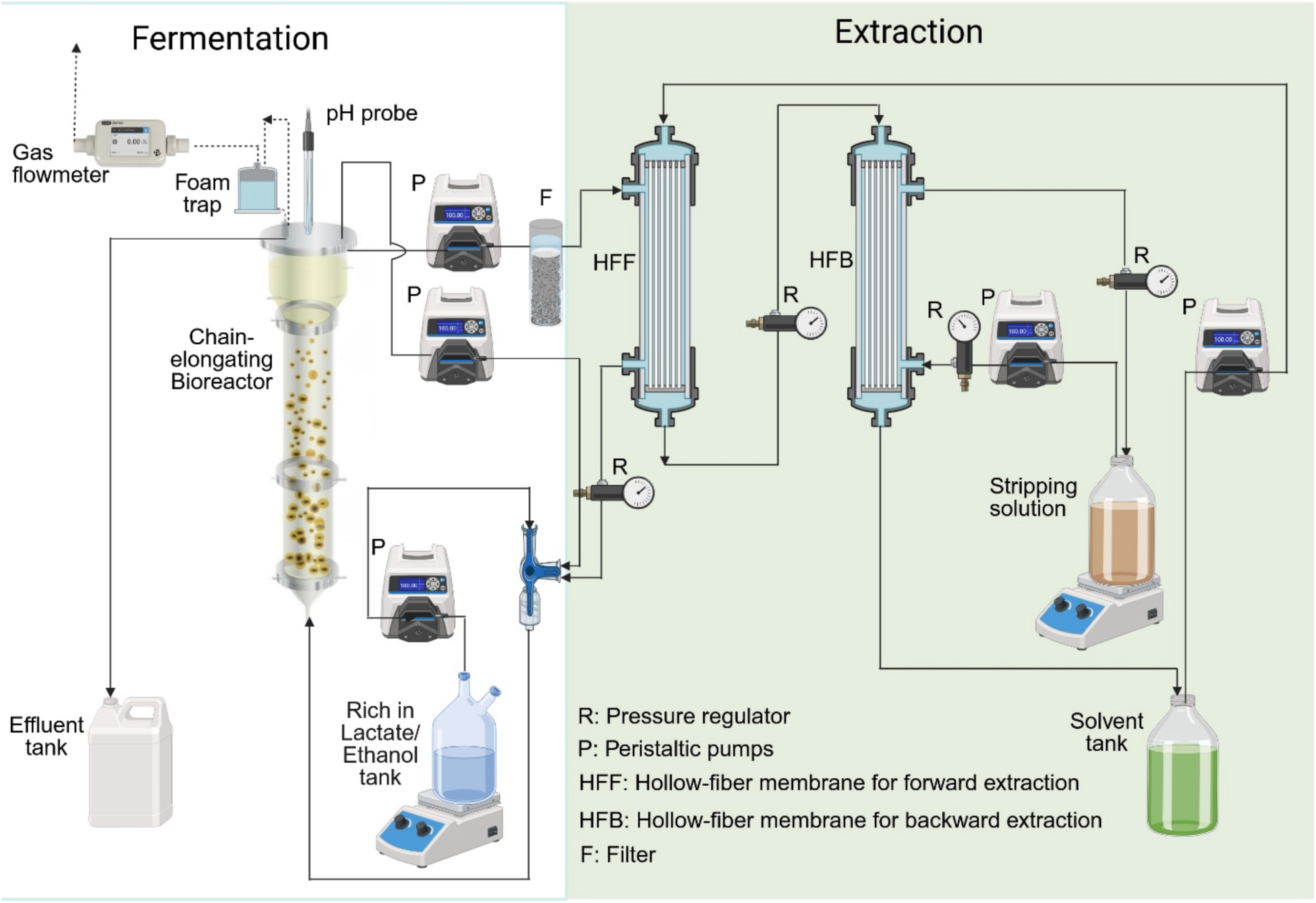
Chain elongation step, including bioconversion and product extraction (step 3 in the overall process flow of the bakery-waste conversion system).

#### 2.2.3. Chain elongation system operating periods

We divided the operating periods into Periods 0-2. An operating period of 92 days (**Period 0**) (data not shown) was initiated to learn how to perform start-up (**Table S2,** ESI †). First, we tested different anaerobic digester sludges to ascertain the preferred inoculum. We compared three different inocula: **(1)** sludge from an anaerobic digester treating manure; **(2)** sludge from an anaerobic digester treating waste-activated sludge; and **(3)** a combination of these two inocula (1:1). Each sludge passed through an 0.6-mm screen mesh to remove large particles. Second, for the combination of the inocula (the preferred inoculum), we determined the ideal operating parameters for the start-up period: hydraulic retention time (HRT); the recycling flow rate of the fermentation broth; the flow direction of the fermentation broth; the hydrophobic solvent in forward and backward membrane contactors; and the substrate concentration to start with (**Table S2,** ESI †).

We indicated the first reported day as Day 1 during **Period 1** (Day 1–44). During this operating period, we wanted to reach a recommended volumetric production rate of caproate >60 mmol C L^-1^ d^-1^ as fast as possible (Kim, B. C. *et al*., 2021). Therefore, we added the electron acceptor *n*-butyrate to the electron donor lactate at final concentrations of 50 mM and 160 mM, respectively. Similarly, acetate was added to ethanol at final concentration of 50 mM and 150 mM, respectively. After we reached the targeted *n*-caproate volumetric production rates, we stopped the external supply of electron acceptors for both bioreactor systems during **Period 2** (Day 44–386). For the lactate medium, we increased the lactate concentration in increments of 25 mM after every 2 HRTs from 160 mM to 260 mM by reducing the dilution with tap water. For the ethanol medium, we used increments of 25 mM to increase the ethanol concentration from 150 mM to 190 mM. During these operating periods, we maintained the systems’ anaerobic conditions because a peristaltic pump connected to a time switch pulsed the reducing agent solution (3M Na_2_S) periodically into the recycling flow of the fermentation broth for each bioreactor system. The pump was on 1 min every 3 h, totaling 8 min d^-1^.

### 2.3. Analytical procedures and calculations

During the pretreatment step, after collecting and drying the bakery waste, we first analyzed the starch content of the dried bakery waste using the rapid total starch (RTS) and the rapid total starch NaOH (RTS-NaOH) method (Total Starch [AA/AMG]) Assay Kit 2020, amyloglucosidase/alpha-amylase method, Neogen, Michigan, US). Then, in the conversion steps, we used high-performance liquid chromatography (HPLC) (Shimadzu, Kyoto, Japan) to measure lactate, ethanol, and simple sugars. We used the total organic carbon (TOC) method (TOC-LCPN FA, E200, Shimadzu, Kyoto, Japan) to measure the carbon content of unmetabolized and unknown compounds. For the chain elongation step, we first collected 2 mL of liquid samples from the bioreactors’ fermentation broth and alkaline extraction solution daily (at the beginning) and twice a week afterwards. The fermentation broth samples were taken from a sampling port at the top of the bioreactors and were divided into two parts: **(a)** for immediate measurement of total ammoniacal nitrogen (*i.e*., both the free ammonia (NH_3_) and ammonium salt (NH_4_^+^) in NH_4_^+^-N); and we calculated the free ammonia based on **Eq. S7** (ESI†); and **(b)** for the measurement of MCCs in effluent. To collect alkaline-extraction solution samples, we provided a T-shaped sampling adaptor in the recirculation line between the reservoir tank exit and the backward membrane contactor. We stored all the liquid samples in 1.5-mL Eppendorf tubes at-20°C until analysis. Next, we collected and immediately measured the gas samples twice a week at the gas sampling port between the top of the bioreactors and the gas flow measuring equipment (Ritter MilliGascounter, Dr.-Ing. Ritter Apparatebau GmbH & Co.KG, Bochum, Germany).

We used an Agilent 7890B gas chromatograph (GC) (Agilent Technologies Inc., Santa Clara, CA, USA) to measure the concentrations of carboxylates (Gemeinhardt *et al*., 2025) and a gas chromatography (GC-FID) (SRI gas GCs, SRI Instruments, Bad Honnef, Germany) for biogas measurement (Gemeinhardt *et al*., 2025). We normalized all given values to mmol carbon atoms (mmol C). We developed a volumetric production rate and specificity equations based on previously reported equations to compensate for the dilution of carboxylates in the alkaline extraction solution (**Eq. S8**, ESI†) (Palomo-Briones *et al*., 2022) and (**Eq. S8,** ESI †) (Xu *et al*., 2018), respectively.

## 3. Results and discussion

### 3.1. Optimized pretreatment conditions were determined for lactate and ethanol production from dried bakery waste

We converted three different dried bakery wastes based on their size into glucose by enzymatic treatment. The conversion efficiency for the small-particle dried bakery waste (≤ 362 μm), medium-particle dried bakery waste (between 360 and 680 μm), and large-particle dried bakery waste (≥ 680 μm) into glucose was 98%±0.66%, 62%±0.36%, and 50%±0.25% (triplicate measurement), respectively (**Fig. S3,** ESI †). The results showed that solids-particle size affected the release of glucose from the bakery waste, which might be due to: **(a)** the particle sizes changing the properties of bakery waste, including the total surface area, and therefore affecting substrate-enzyme interaction (Guerra-Oliveira *et al*., 2022); **(b)** the increasing solid content hindering the substrate-enzyme interaction (Thani *et al*., 2021); **(c)** the protein matrix affecting the shape and accessibility of starch granules; and **(d)** the increasing protein content resulting in the deformation of the starch structure, making it difficult for the starch to be degraded (Agu *et al*., 2009). The results showed that the dried bakery waste with small particle sizes is superior. However, small particle sizes could increase the downstream load of the centrifuge. Therefore, the choice of particle size must be a compromise between yield and minimizing problems for the downstream process.

The thermodynamic capacities of lactate and ethanol to operate as electron donors for chain elongation are similar (Weimer *et al*., 2016). Lactate conversion to MCCs releases CO_2_, while ethanol conversion does not, because ethanol has an average carbon atom oxidation number that is more reduced than that of lactate. Because of similar thermodynamic capacities, we optimized the conversion of the glucose-rich solution into both lactate and ethanol (**Table S3**, ESI †). For lactate production, the results showed that the small-particle mash that was pretreated with *B. coagulans* DSM1 produced the highest lactate concentration (263±0.120 mmol L^-1^) at a temperature of 50^°^C compared to the other inoculants (**Table S3**, ESI †). In addition, the large-particle mash that was pretreated with the lactate microbiome performed better than *B. coagulans* and Silosolve MC at low temperatures. For example, at 25^°^C, the lactate concentrations were 121±41.1, 55.1±10.2, and 45.2 ± 14.2 mmol L^-1^ using the lactate microbiome, *B. coagulans* DSM1, and Silosolve MC, respectively (**Table S3**, ESI †). This might be explained by the fact that the lactate microbiome was an open culture with different microbial populations, which could be active at different temperatures (Schuetterle *et al*., 2024). The disadvantage of using a lactate microbiome was that it produced a mixture of lactate and ethanol, reducing the selectivity for lactate production. Silosolve MC performed worst as an inoculant because it is intended to control fermentation during silage (Daniel *et al*., 2018). For ethanol, the maximum concentration was 180 ± 0.22 mmol L^-1^ at 30°C, which is the optimum temperature for technical yeast. In general, we observed the same advantage of the smaller particle size in mashes for the lactate and ethanol production compared to the glucose production: small-particle mash > medium-particle mash > large-particle mash (**Fig. S3, Table S3,** ESI †). Therefore, for the rest of the microbial chain elongation study, we decided to use: **(a)** the small-particle fermented mash; **(b)** the *B. coagulans* DSM1 inoculum to produce lactate; and **(c)** the technical yeast to produce ethanol.

### 3.2. The bakery-waste solids reduced the raffinate flux for the extraction system that will be coupled to the chain elongation system

As discussed in the introduction, starch-rich substrates can cause stickiness. In addition, particles from bakery waste can also form cakes. Therefore, we performed short-term experiments (24 h) to test how fouling of our hollow-fiber membranes throughout longer operating periods could negatively impact pertraction. Under normal operating conditions, the undissociated *n*-caproic acid diffuses from the fermentation broth to the solvent, and thus no liquid is pushed through the membrane with small holes of around 200 nm. Our 24-h experiment was divided into two sections. First, we fouled the membrane intentionally by pushing the various fermented mash types through our membrane at a constant pump rate for 12 h during which we studied the decline in raffinate flux (*i.e.*, the clear liquid). We observed for all fermented mash types that the decline in raffinate flux can be divided into two decay periods: a sharp-decay period (0–2 h) and a pseudo-steady-state-decay period (2–12 h) (**Fig. 2A**). The reduction in raffinate flux was caused by a constant increase in solids retardation, depositing a cake/gel layer on the hollow-fiber membrane, especially for the large-particle fermented mash (**Fig. 2B**). For the small-particle fermented mash, we still observed a sharp-decay period, albeit not as dramatic as the medium-and large-particle fermented mash types (**Fig. 2A**), even though the fouling rate was minimal. We do not know why the raffinate flux decreased sharply during the sharp-decay period for the small-particle fermented mash Second, we performed conventional pertraction for 12 h using the hollow-fiber membranes from the three fermented mash types after 12 h during the fouling section. The increasing amounts of formed cake/gel on the surface of the hollow-fiber membranes from the small-, medium, and large-particle fermented mash resulted in an increased resistance during pertraction, leading to a slower extraction rate of undissociated *n*-caproic acid throughout the 12-h experimental period of section 2 (**Fig. 2C**). De Sitter *et al*. (2018) found the same observation with different hollow-fiber membrane properties. The effect of bakery-waste solids on the extraction system showed that the small-particle mash performed better than medium-particle and large-particle mashes (**Fig. 2C**). The extraction of *n*-caproate was reduced when the fouling factor of the membrane increased; for example, we extracted 40 mM and 15 mM when the fouling factor was 4×10^-3^ mg L^-1^ m^-2^ and 3.75 mg L^-1^ m^-2^, respectively (**Fig. 2C**). Therefore, we used small-particle bakery-waste mash for the chain elongation experiments to avoid excessive fouling of the membrane for MCC extraction.

**Figure 2.**
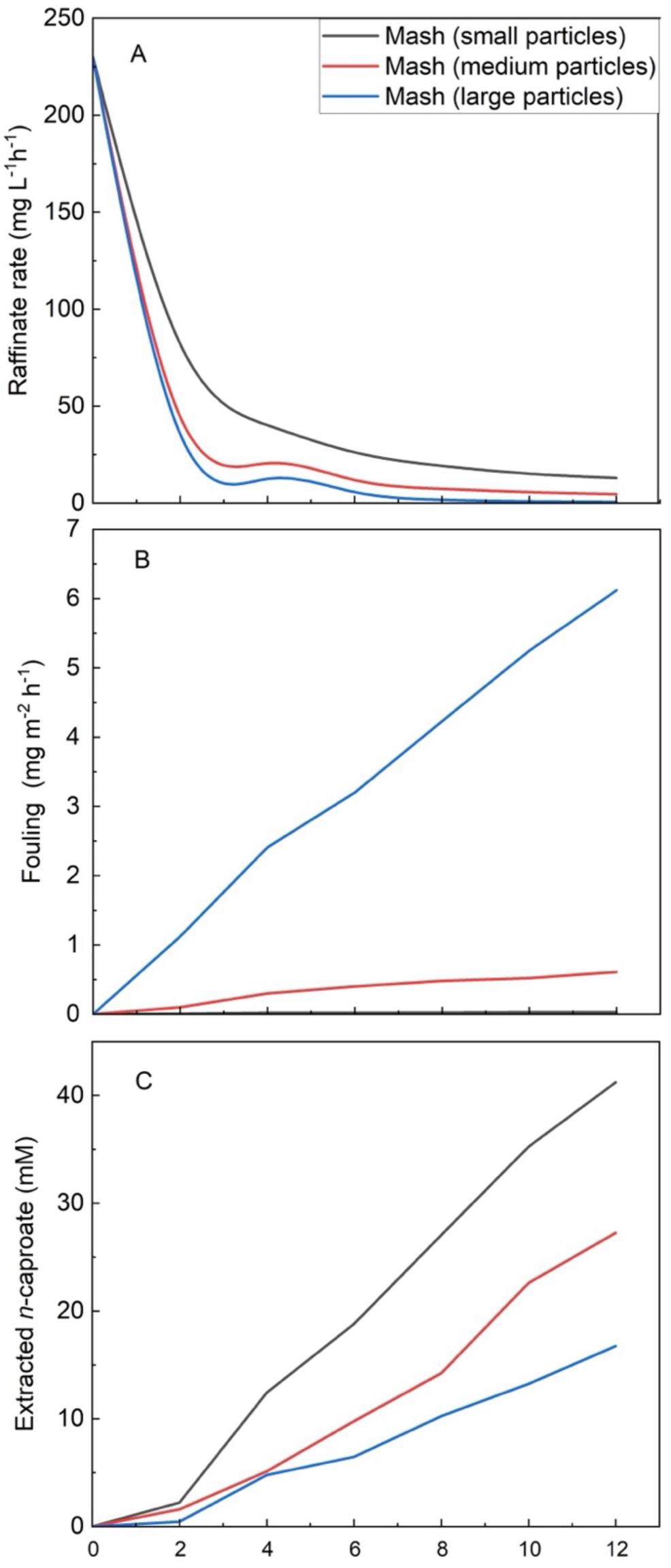
Effect of different fermented mash types on fouling and MCC extraction for the hollow fiber-membrane: **A**) raffinate rate during the 12-h fouling experiment, **B**) fouling effect during the 12-h fouling experiment, and **C**) extracted *n*-caproate during the 12-h extraction experiment.

### 3.3. Only the chain elongation experiment with the intermediate lactate as the electron donor resulted in satisfactory MCC selectivities

Initially, we intended to use and compare lactate-rich and ethanol-rich substrates to produce MCCs. Unfortunately, the bioreactor fed with an ethanol-rich substrate did not behave as envisioned and produced excessive methane, which is typical for an anaerobic digester. We were unable to improve performance, even during a lengthy operating period, despite implementing several operating optimizations. Possibly, the ratio of ethanol (electron donor) to other substrates (electron acceptor) was too low to produce MCCs, lacking enough reducing power. Therefore, we summarized the results of the bioreactor fed with the ethanol-rich substrate in the supporting information (**Result summary of ethanol-bioreactor, Fig. S4-S5,** ESI †). Here, we focused on the performance and behavior of the chain-elongating bioreactor that was fed with a lactate-rich substrate.

#### 3.3.1. Chain elongation with lactate as the electron donor, with an external *n*-butyrate supply, was accelerated

A start-up study with reactor microbiomes achieved increased volumetric *n*-caproate production rates during a short operating period after inoculation with anaerobic digester sludge. They added *n*-butyrate as an electron acceptor, in addition to the electron donor, for chain elongation of different substrates (Kim, B. C. *et al*., 2021; Nzeteu *et al*., 2022). Acetate as an alternative electron acceptor would have been a possibility, but Wu *et al*. (2019) showed a low *n*-caproate volumetric production with acetate. Therefore, we shaped the *n*-caproate-producing microbiome by using lactate as the electron donor and by externally adding *n*-butyrate as an electron acceptor, and utilizing a mixed inoculum (**The choice of inoculum,** ESI †). We know from previous studies that feeding primarily electron donors, such as lactate and ethanol, is also possible, but that a lack of growth must be anticipated (Allaart *et al*., 2024). Therefore, we opted to externally add electron acceptors to speed up the growth and acclimation of the microbiome during the initial start-up period for the chain-elongating bioreactors.

From Day 1 to 44, the volumetric production rates of *n*-caproate and *n*-caprylate increased steadily, reaching 70 mmol C L^-1^d^-1^ and 18 mmol C L^-1^d^-1^, respectively (**Fig. 3A**). For *n*-Butyrate, we found a fluctuating volumetric production rate of 38 mmol C L^-1^d^-1^ and 50 mmol C L^-1^d^-1^, while the volumetric production rate for acetate reached 22 mmol C L^-1^d^-1^ (**Fig. 3A**). In addition, the carbon mass balance showed that 30.6% of the carbon fed in the upflow bioreactor ended up in CO_2_ production (**Fig. 3B**), which is as anticipated because one carbon atom out of three carbon atoms of lactate is oxidized to CO_2_, according to the stoichiometry. Kim, B. C. *et al*. (2021) showed that the addition of *n*-butyrate to lactate as an electron donor quickly (84 h) acclimated the MCC-producing microbiome during bottle experiments (batch-fed, non-open culture). We also found that in a continuously fed open-culture bioreactor, the enrichment of an MCC-producing microbiome progressed rapidly (a couple of days in (**Fig. 3**)).

**Figure 3.**
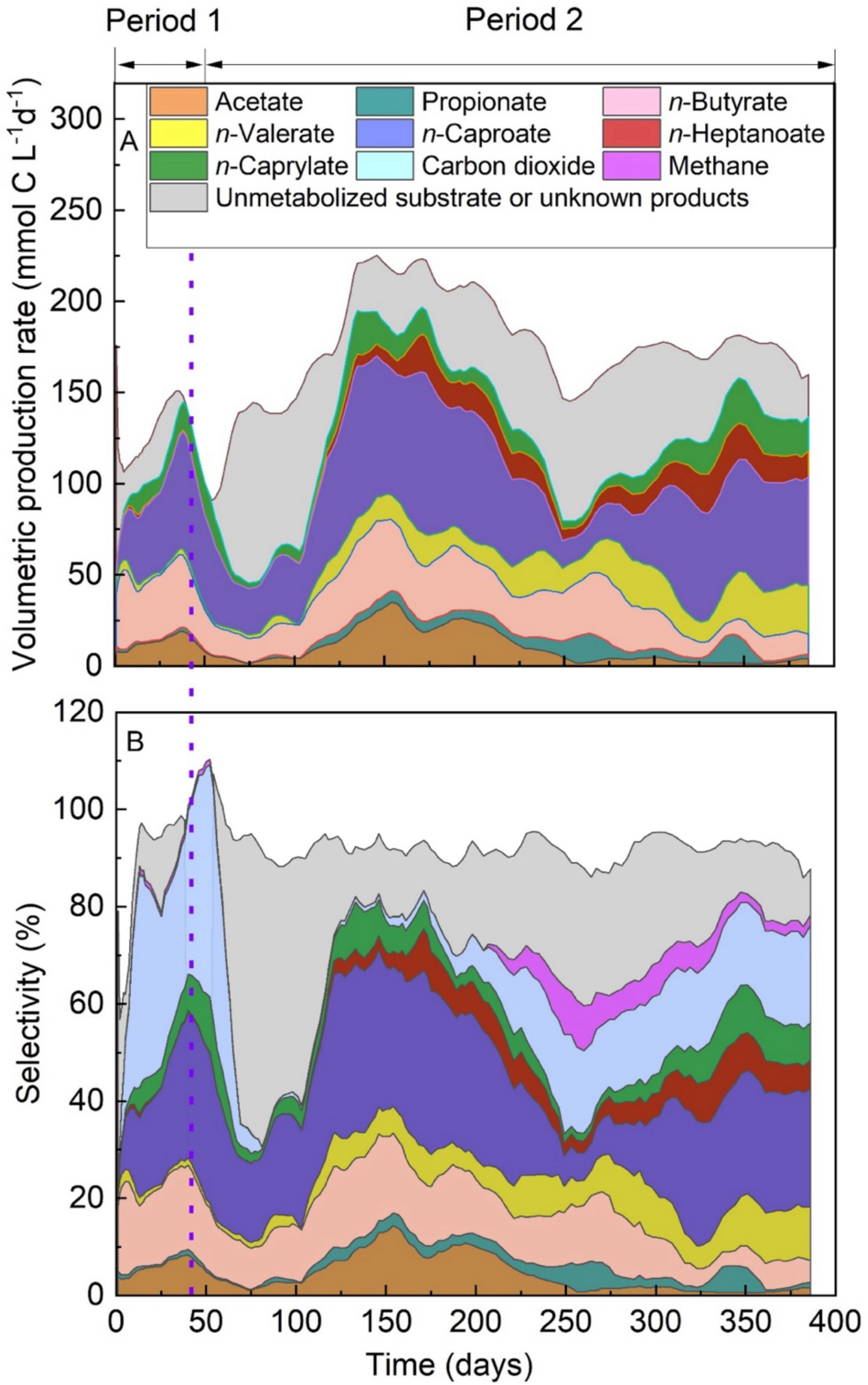
Carboxylate, unmetabolized substrate/unknown product, and gas profiles throughout the operating period from the bioreactor that was fed with lactate as an intermediate after pretreatment of bakery waste: **A**) volumetric production rates, and **B**) selectivity. The volumetric production rates and selectivities were calculated, and the graphs were smoothed with a six-measurement moving average. Operating periods 1 and 2 are indicated and described in the Materials and Methods section.

#### 3.3.2. More research is necessary to understand whether the external addition of *n*-butryate actually shortened the microbiome acclimation period or not

However, after 44 days, when we removed the externally added *n*-butyrate, because we wanted to only use lactate as the substrate, unstable conditions occurred (broken line on Day 44 in **Fig. 3**). The absence of *n*-butyrate led to a large amount of unmetabolized substrate and unknown products (enlarged gray area in **Fig. 3**), and a decrease in the volumetric *n*-caproate production rate from 64.76 to 23.68 mmol C L^-1^d^-1^ on Day 44 and 90, respectively (**Fig. 3A**). Due to these unstable conditions, we needed an additional acclimation period (**Fig. 3**).

Between Days 44 and 129, we observed an increase in the volumetric production rate of odd-number carboxylates, especially *n*-valerate and *n*-heptanoate, which increased 6-fold (**Fig. 3A**). We also observed a gradual increase in the volumetric *n*-caproate production rates from 64.76 mmol C L^-1^d^-1^ on Day 44 to 81.68 mmol C L^-1^d^-1^ on Day 129 (**Fig. 3A**). Thus, the combined acclimation periods took 129 days.

We hypothesized that circumventing *n*-butyrate led to: **(1**) the disruption of redox balance, leading to a microbial metabolic shift, which caused the production of undesired products and unmetabolized substrate, which resulted in a decrease in SCC production; and (**2**) the degradation of protein into amino acids, contributing to odd-number carboxylate production. Steinbusch *et al*. (2011) state that internal redox balancing and microbial metabolism stability are necessary for the chain elongation process without an electron acceptor. Here, it took approximately another 90 days for redox balancing and microbial metabolism to stabilize after removing the external electron acceptor (**Fig. 3B**). We observed that between Day 75 and Day 175, the specificity of CO_2_ was close to zero (**Fig. 3B**). This operating period included the acclimating period during which redox balancing must have occurred. We observed an increase in homoacetogenesis (*i.e.*, acetate production) during this period with an excess of reducing equivalents due to the sudden absence of the external electron acceptor from *n-*butyrate, because methane production was still absent (**Fig. 3B**). Indeed, the selectivity of acetate went from 2% to 16% between Day 75 and Day 175 (**Fig. 3B**), explaining why we did not observe CO_2_ in headspace as anticipated by the stoichiometry.

The maximum volumetric *n*-caproate production rate of 94.2 mmol C L^-1^d^-1^ (0.08 g L^-1^ h^-1^) was achieved on Day 175 of the operating period (**Fig. 4A**), which coincided with the maximum *n*-caproate selectivity of 55% (**Fig. 3B**). The acclimation of the microbiome continued with the highest volumetric *n*-valerate (unwanted) and *n*-caprylate (wanted) of 42.8 mmol C L^-1^d^-1^ and 26.8 mmol C L^-1^d^-1^ being achieved on Day 235 and Day 351, respectively (**Fig. 4A**). A continuous reduction in the performance between Day 175 and Day 248 was due to a faulty pH probe, and the system recovered again after this experimental problem had been mitigated (**Fig. 4A**). A stable maximum average *n*-caprylate specificity of 11.1 ± 6.88% (n=387) was achieved at the end of the operating period during which the maximum average overall MCC specificity was 55.2 ± 14.1 % (n=386; **Fig. 4A**). We would have to perform a side-by-side study to investigate whether the external addition of *n*-butyrate, which looked promising during the first 44 days of the operating period (**Fig. 3B**), compared to the absence of *n*-butyrate would resulted in a different total acclimation period for the microbiome to produce MCCs.

**Figure 4.**
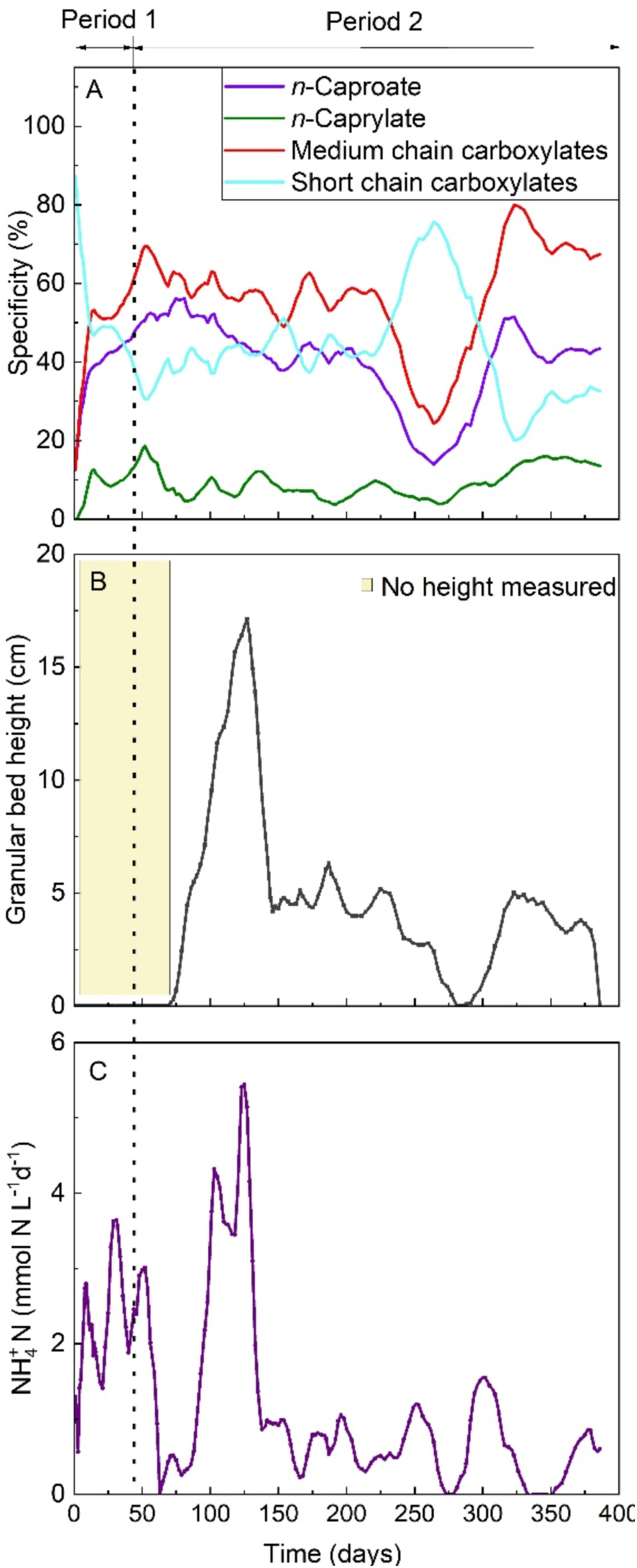
Effect of pseudo-granules and ammonium nitrogen concentrations on the specificities of carboxylates: **A**) specificity of carboxylates, **B**) pseudo-granular bed height in the bioreactor, and **C**) volumetric ammonium nitrogen production rate. The product specificities, pseudo-granular bed heights, and volumetric ammonium nitrogen production rates were calculated, and the graphs were smoothed with a six-measurement moving average.

#### 3.3.3. The lactate-fed chain elongation bioreactor formed pseudo-granules, which were not stable

During Period 1, we observed the presence of a few pseudo-granules at the bottom of the bioreactor; however, we had not yet measured the granular bed height in the bioreactor. At the beginning of Period 2, the pseudo-granules grew in numbers, and we started to measure the granular bed height (**Fig. 4B**). The granules were not stable, and we observed several rapid drops in granular bed height, including on Day 133 of the operating period (**Fig. 4B**). This change occurred without changes in the performance (**Fig. 3A-B**), which is why we called them pseudo-granules (**Fig. S6**, ESI †). We were unable to remove the pseudo-granules from the bioreactor to analyze and confirm their nature. Real anaerobic granules remain as individual entities even after sludge settling; however, we could not perform this measurement without disintegrating the pseudo-granules. According to Roghair *et al*. (2016), chain elongation with granular sludge could become a high-rate biotechnological process. However, based on the disintegration of our pseudo-granules, developing a granulation concept around the chain elongation of the dried-bakery waste would not be feasible. In addition, the presence of pseudo-granules had a little impact on volumetric production rate and specificity for *n*-caproate and *n*-caprylate (**Fig. 3A, Fig. 4B**).

#### 3.3.4. Ammonium-nitrogen production from protein degradation is pertinent as a nitrogen source and pH control

The volumetric ammonium-N production rate increased to 2.8 mmol N L^-1^ d^-1^ from Day 0 to Day 18 of the operating period, and then fluctuated between 2.8 mmol NL^-1^d^-1^ and 3.9 mmol N L^-1^ d^-1^ until Day 44 (**Fig. 4C**). A lower biological activity after removing the external *n*-butyrate addition on Day 44 resulted in a considerably reduced volumetric ammonium-N production rate (**Fig. 4C**). The acclimation of biomass due to pseudo-granulation occurred after the removal of external *n*-butyrate, and was likely not due to the changes in ammonium-N concentrations. The pseudo-granulation on Day 74 started before a renewed sharp increase in the ammonium-N concentration on Day 83. In addition, during Period I, no pseudo-granules were observed even though a relatively high concentration of ammonium-N was achieved (**Fig. 4B**, **Fig. 4C**).

From Day 62 to Day 125, the volumetric ammonium-N production rate increased sharply with a maximum of 5.6 mmol N L^-1^ d^-1^ on Day 127 of the operating period (**Fig. 4C**). We observed a sharp decline after which the volumetric ammonium-N production rate fluctuated, but remained below 1 mmol N L^-1^ d^-1^ until Day 386 (**Fig 4C**). We do not understand why we observed these fluctuations. We speculate an interplay between the following: **(1)** protein decomposition to produce ammonium-N; **(2)** fluctuation of the incoming protein concentration in the lactate-rich substrate; **(3)** fluctuations in operating parameters (*e.g.*, cell optical density, pH, and temperature); and **(4)** removal of ammonium-N due to biomass growth. We also observed a variation in the concentration of ammonium-N along the effective height of the bioreactor (data not shown). The latter variation could be the result of varying biomass concentrations along the effective height of the bioreactor. However, the fluctuations of the volumetric ammonium-N production rates had little impact on the volumetric production rate of *n*-caproate and *n*-caprylate (**Fig. 3A, Fig. 4C**).

#### 3.3.5. Sankey diagram for bakery waste conversion *via* microbial chain elongation

The Sankey diagram with the carbon flow in our bakery-waste fermentation system provides a summary and clear visual representation of microbial chain elongation with (**Fig. 5A**) and without (**Fig. 5B**) *n*-butyrate as the external electron acceptor. We normalized all measured values of TOC in bakery waste to 100 kg of carbon atoms (kg C), which represented 143 kg of dried bakery waste. The microbial chain-elongation of bakery waste with *n*-butyrate as an externally added eletron acceptor resulted in the wanted products of 20.1 kg C of *n*-caproate and 4.74 kg C of *n*-caprylate, while the co-products consisted of: 0.490 kg C of *n*-heptanoate; 26.5 kg C of SCCs; 32.0 kg C of CO_2_; 0.977 kg C of CH_4_; and 32.9 kg C of unmetabolized substrate or unknown products (**Fig. 5A**). The microbial chain elongation without *n*-butyrate addition resulted in 22.1 kg C of *n*-caproate and 5.00 kg C of *n*-caprylate as wanted product, which is slightly higher but not that different from when *n*-butyrate was added. However, in this representation, which was based on 100 kg of bakery waste, we did not account for the kg of C in the added *n*-butyrate. For without *n*-butyrate, the co-products were: 4.20 kg C of *n*-heptanoate; 23.1 kg C of SCCs, 10.1 kg C of CO_2_, 2.21 kg C of CH_4_, and 32.4 kg C of unmetabolized substrate or unknown products (**Fig. 5B**). The difference between with and without *n*-butyrate was mostly in the reduced selectivity of CO_2_. We hypothesize that without *n*-butyrate, the microbiome is forced to produce its own electron acceptor, acetate, through acetogenesis, thereby reducing the amount of CO_2_ in the bioreactor’s headspace.

**Figure 5.**
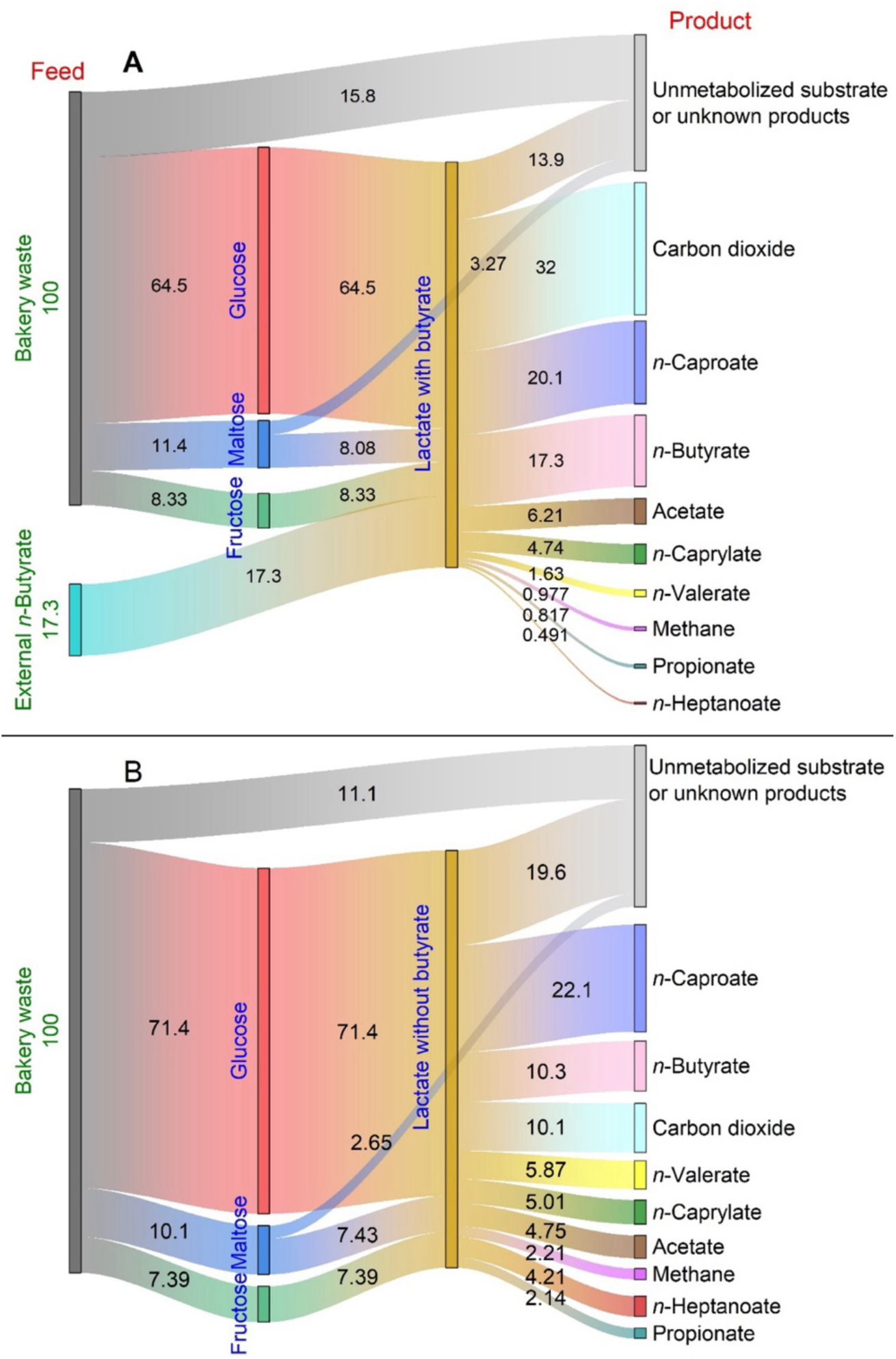
Sankey diagram for bakery-waste conversion *via* pretreatment steps and microbial chain elongation: **A**) chain elongation with the supply of external *n*-butyrate, and **B**) chain elongation without the supply of external *n*-butyrate (unit is kg C).

The carbon of unmetabolized or unknown products included the carbon lost during the size reduction in the bakery waste. Clearly, there is a need for improvement to recover residual carbon (unmetabolized substrate and unknown products) (**Fig. 5A**, **Fig. 5B**). Nevertheless, we achieved a higher carbon conversion to *n*-caproate and *n*-caprylate compared to food wastes in other studies. For example, Huo *et al*. (2024) in their chain elongation study, the conversion from food waste carbon to *n*-caproate was 15.5%, with no carbon converted to *n*-caprylate. Therefore, our study is among the few studies that have converted food waste that is rich in carbohydrates into *n*-caprylate using continuous chain elongation coupled with pertraction.

## 4. Conclusions

We investigated the recovery of carbon in bakery waste by first converting it into the intermediate lactate, which was then utilized as an electron donor for the microbial chain elongation process to produce medium-chain carboxylates. We achieved a satisfactory medium-chain carboxylate production rate for lactate as an electron donor. However, our efforts with ethanol as an intermediate did not result in a satisfactory production rate. Our results identified that the chain elongation process with lactate, when coupled to pertraction, required: **(a)** the filtration of bakery-waste mash to avoid fouling of the membrane, consequently optimizing pertraction; **(b)** a mash containing solid particles with an average size distribution of ≤ 362 μm; and **(c)** *Bacillus coagulans* DSM1 inoculum to produce lactate effectively. The bakery waste mash with small particle sizes was the best for pretreatment and pertraction. However, small particle sizes might increase the centrifuge load in large-scale applications. Thus, the choice of particle size must be a compromise between yield and minimizing problems in the downstream process. The results of this study inspire the development of bakery waste pretreatment, chain elongation, and product recovery strategies for *n*-caproate and *n*-caprylate production, as well as the conversion of the processing system’s residual solids into methane. We refer to this process as the bakaroate process.

## CRediT authorship contribution statement

**Jean Népomuscène Ntihuga:** Conceptualization, data curation, formal analysis, investigation, methodology, project administration, visualization, writing-origin draft, writing-review & editing. **Joseph G. Usack:** Conceptualization, project administration, supervision, writing-review & editing, **Rafael Mayer:** data curation, writing-review & editing. **Ahmed K. Kinbokun:** data curation, writing-review & editing. **Hatice Yasil:** data curation, writing-review & editing. **Mei Zhou:** data curation, writing-review & editing. **Largus T Angenent:** Conceptualization, funding acquisition, project administration, resources, supervision, and writing-review & editing

## Acknowledgments

Largus T. Angenent acknowledges the support from the Alexander von Humboldt Foundation within the framework of the Alexander von Humboldt Professorship, which is endowed by the Federal Ministry of Education and Research in Germany. The authors acknowledge the support of Bäckerhaus Veit for providing the bakery waste at no cost. We thank Abha Mansoor and Fady Isaac from the University of Tübingen for their support in operating the bioreactors. Finally, we thank a group of M.Sc. students, consisting of Chenhui Chang, Chih-Fu Cheng, and Elena Schiller, for their contribution to the ideation process.

