## Supporting Information for "Dried-bakery waste as a substrate for *n*-caproate and *n*-caprylate production *via* chain elongation: bakeroate"

### Experimental procedures

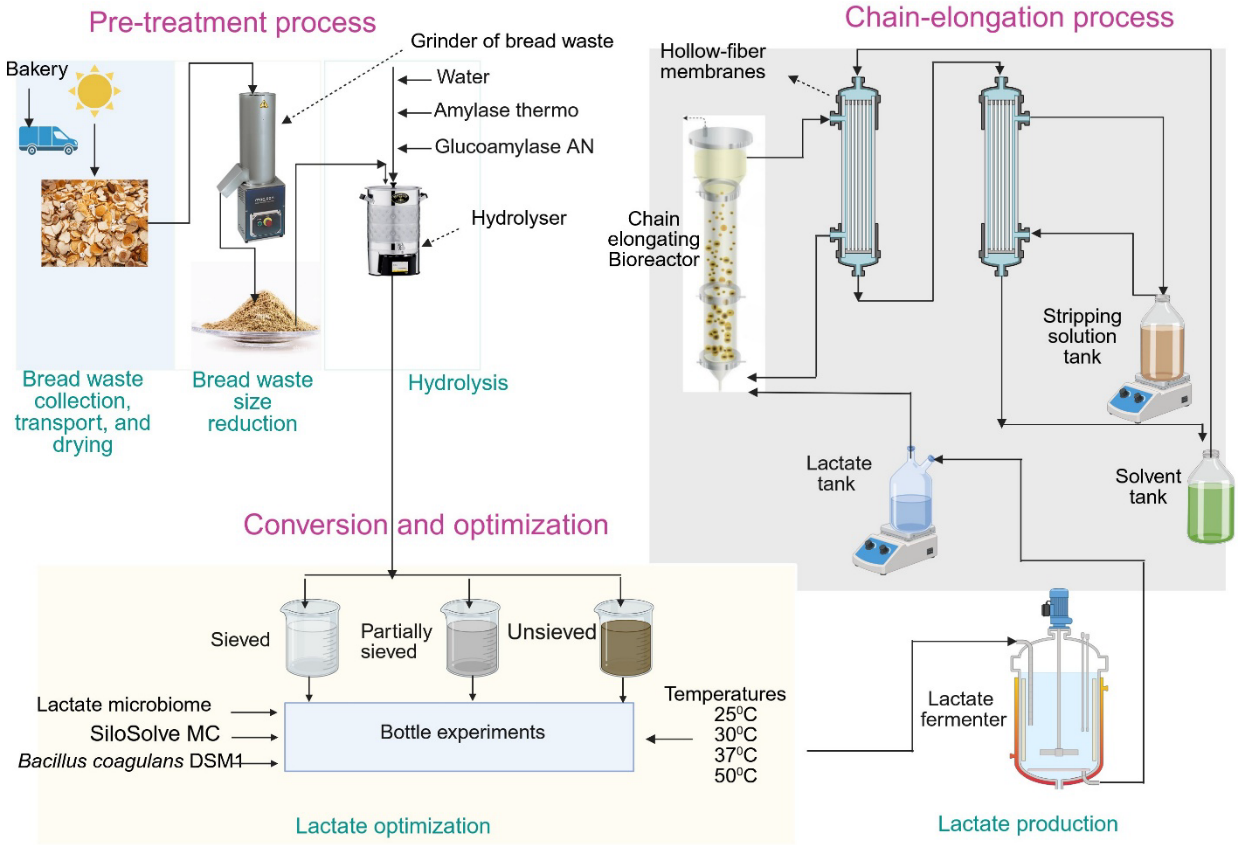


**Figure S1**. Overall process flow for bakery-waste conversion, utilizing lactate as an intermediate.

**Table S1.** Bakery-waste composition (data from Bäckerhaus Veit).

| Name | Composition in 100 g |
|---|---|
| Fat | 1.40 |
| Saturated fatty acid | 0.40 |
| Carbohydrates | 54.0 |
| Added sugar | 1.00 |
| Dietary fiber (non-starch polysaccharides) | 3.10 |
| Protein | 9.20 |
| Salt | 1.60 |

**Tryptone soya medium**

1.7% (w/v) pancreatic digest of casein, 0.3% (w/v) soybean meal, 0.5% (w/v) NaCl, 0.25% (w/v), $KH_2PO_4$, 0.25% (w/v), glucose].

**Lactate and ethanol preparations**

We prepared the medium stock in a 10-L bottle by first adding 8 L lactate or ethanol (diluted if necessary) and then successively adding 100 mL trace metal stock solution, 12.5 g yeast extract, 30 g sodium bicarbonate, 50 g MES (2-morpholinoethanesulphonic acid), and 49 mL *n*-butyrate for the lactate substrate and 14.9 mL acetate for the ethanol substrate. Afterwards, we filled the mixture to 10 L with lactate or ethanol mashes and shook the bottle in rotary movements to homogenize the content. We adjusted the pH between 5.0 and 5.4 using sodium hydroxide or hydrochloric acid. The medium preparation was performed under ambient air without sterilization, and the 10-L medium stock beaker was kept in a 4°C storage room until usage. When the bioreactor system's 5-L medium supply tanks ran out of medium, we refilled them with fresh medium from the medium stock.

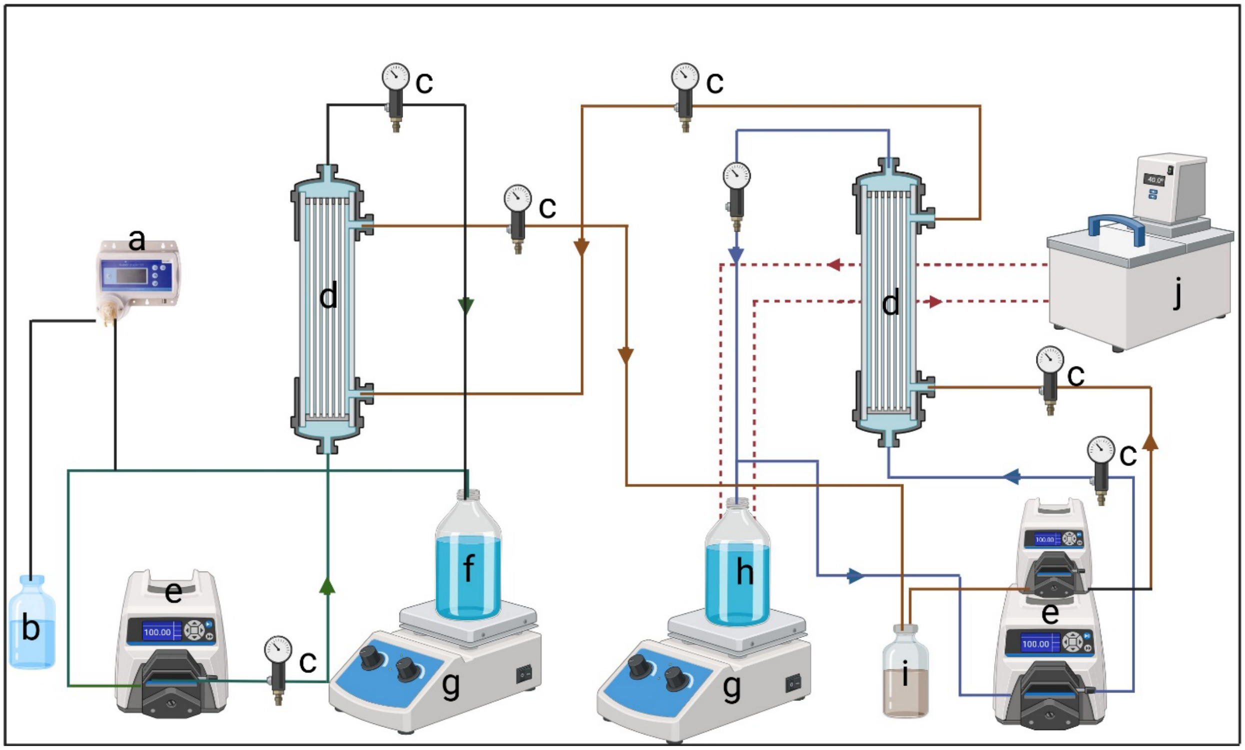


**Figure S2**. Fouling and extraction experiment set-up: a) pH controller; b) NaOH tank; c) pressure meters; d) hollow-fiber membranes; e) pumps; f) stripping tank; g) mixers; h) mash tank; i) solvent tank; and j) water bath.

### Batch extraction calculations

$t$ = The duration of the batch experiment (h), and i = 2 to 12

$W_{ref\_i}$ = Reference weight of the solvent, g

$W_{S\_i}$ = Final product weight after the batch extraction, g

$W_{EC6\_i}$ = Weight of extracted *n*-caproic acid, g

$$W_{EC6\_i} = W_{S\_i} - W_{ref\_i} \text{ , g}$$

**Description of the extraction system**

We modified the pertraction system as reported previously to extract MCCs (Agler *et al.*, 2012; Ge *et al.*, 2015). The extraction system consisted of a forward (3M™ Liqui-Cel™ EXF 4X13 Series Membrane Contactor G574, Polypropylene X50, Charlotte, NC, USA) and a backward hollow-fiber membrane contactor (3M™ Liqui-Cel™ EXF 2.5X8 Series Membrane Contactor G648, Polypropylene X50) with a surface area of 8.1 $m^2$ and 1.4 $m^2$ for forward and backward contactors, respectively (**Fig. 1, Fig. S1**). We maintained the temperature of each membrane contactor at 30°C by covering it with a heating mat (Heat mat comfort, Terra Exotica, Alfeld, Germany) that was connected to a temperature controller (Ink bird ITC-308 Temperature Controller, Guangdong, China). A peristaltic pump (Model 7522-30, Cole-Parmer) recycled the fermentation broth between the bioreactor vessel and the forward membrane contactor through tubing (Tygon® A-70-F, Masterflex, Gelsenkirchen, Germany) at a rate of ~6.48 L $h^{-1}$.

A filter module with a wet volume of 0.4 L was connected in-line between the bioreactor vessel and the extraction system to prevent fouling and associated plugging of the hollow-fiber membranes for the first 186 days of the operating period. It was later replaced with a 0.8-L module (GXWH20S, FXWSC filter, General Electric Company, Boston, MA, USA). The hydrophobic solvent, which comprised of mineral oil with 30 g $L^{-1}$ trioctylphosphine oxide (TOPO), was recycled with a peristaltic pump (Model 7522-30, Cole Parmer) through the hollow-fiber lumen side of the forward and shell side of the backward contactor at a flow rate of 4.32 L $h^{-1}$ in both systems. An alkaline extraction solution, which was a mixture of sodium borate (30 g $L^{-1}$) and sodium hydroxide (7 g $L^{-1}$), was recycled with a pump (Model 07528-30, Cole Parmer) between the lumen side of the backward contactor and a 5-L stirred vessel at a flow rate of 4.32 L $h^{-1}$ in both bioreactor systems. A pH electrode (ProcessLine 80-325 pH, SI Analytics, Weilheim, Germany), which was attached to a pH controller (Bluelab pH controller, Bluelab Corporation Limited, Tauranga, New Zealand), automatically adjusted the pH of the alkaline extraction solution to a pH of 9 with sodium hydroxide (5 M).

**Table S2.** Experimental parameters.

| Name | Lactate bioreactor | Ethanol bioreactor |
|---|---|---|
| Main substrates < 50 days | Lactate and butyrate | Ethanol and acetate |
| Main substrate > 50 days | Lactate | Ethanol |
| Operating temperatures (°C) | 37.0 | 30.0 |
| Bioreactor mean cross-sectional area ($cm^2$) | 87.1 | 87.1 |
| Bioreactor volume (L) | 6.70 | 6.70 |
| Hydraulic retention time (d) | 3.00 | 3.00 |
| Up-flow velocity ($m\ h^{-1}$) | 3.00 | 3.00 |
| Feeding flow rate ($L\ h^{-1}$) | 0.093 | 0.093 |
| Raffinate flow rate ($L\ h^{-1}$) | 6.48 | 6.48 |
| Biomass flow rate ($L\ h^{-1}$) | 3.52 | 3.52 |
| Direct recirculation rate ($L\ h^{-1}$) | 2.90 | 2.90 |
| Solvent flow rate ($L\ h^{-1}$) | 4.32 | 4.32 |

### Bioreactor medium preparation

The medium was produced in a 10-L beaker. First, we added 8 L of diluted or undiluted (depending on the desired concentration) lactate-rich solution and ethanol-rich solution for the lactate bioreactor system and the ethanol bioreactor system, respectively; we added

- 100 mL trace metal stock solution
- 30 g sodium bicarbonate
- 50 g MES
- 15 g yeast extract

We shook the beaker clockwise and counterclockwise till we had a homogeneous solution. Then, we filled the beaker up to 10 L with diluted or undiluted lactate-rich or ethanol-rich solution. Finally, we adjusted the pH between pH = 5.0 and pH = 5.4 with NaOH or HCl.

### Trace metal stock (g $L^{-1}$)

2 g nitrilotriacetic acid; 1 g manganese sulfate; 0.8 g ammonium iron (II) sulfate; 0.2 g cobalt chloride; 0.2 g zinc sulfate; 0.02 g copper (II) chloride; 0.02 g nickel chloride; 0.02 g sodium molybdate; 0.02 g sodium selenate; 0.02 g sodium tungstate.

### Bioreactor equations

Both bioreactor systems' wet working volumes (L) were the same. The fermentation broth levels and wet working volumes rose after connecting the swan necks to the bioreactor vessels.

The wetted volume of the bioreactor with a filter (0.4 L) and with a 0.8 L

$$V_{wet,x} = V_{Vessel\ with\ swan\ neck} + V_{Filter\ module} \quad \textbf{(Eq. S1)}$$

$V_{Vessel}$ = 5 L  $V_{Filter\ module}$ = 0.4/0.8 L  $V_{vessel\ with\ swan\ neck}$ = 6.7 L

Where:

$V_{Vessel}$ = Wet volume of the bioreactor vessel, L

$V_{Filter\ module}$ = Wet volume of the filter module, L $V_{vessel\ with\ swansneck}$ = Elevated wet volume of the bioreactor vessel with attached swan neck, L

**Total organic loading rate (mmol C L$^{-1}$ d$^{-1}$).**

We calculated the total organic loading rate ($OLR$) based on the content of the influent and converted it into mmol C. The lactate-rich solution included lactate, maltose, glucose, and ethanol, as well as with/or without butyrate. The ethanol-rich solution included ethanol, maltose, glucose, and, with /or without acetate

$$OLR_{Lactate} = \frac{(3xC_{lactate}+12xC_{maltose}+6xC_{glucose}+2xC_{Ethanol}+4xC_{butyrate})\times f}{V_{wet,x}} \quad \textbf{(Eq. S2)}$$

$$OLR_{ethanol} = \frac{(2xC_{ethanol}+12xC_{maltose}+6xC_{glucose}+2xC_{acetate})\times f}{V_{wet,x}} \quad \textbf{(Eq. S3)}$$

Where:

$C_{lactate}$ = Concentration of lactate in the influent, mM

$C_{maltose}$ = Concentration of maltose in the influent, mM

$C_{glucose}$ = Concentration of glucose in the influent, mM

$C_{butyrate}$ = Concentration of *n*-butyrate in the influent, mM

$C_{Ethanol}$ = Concentration of ethanol in the influent, mM

$C_{Acetate}$ = Concentration of acetate in the influent, mM

$f$ = Influent flow rate, L d$^{-1}$

$V_{wet,x}$ = Wet working volume of the system, $x = 0.4\ or\ 0.8$ L

Alkaline extraction solution volume on day n (L). To account for the dilution of accumulating carboxylates, we extrapolated the volume of the alkaline extraction solution on day n ($V_{ex,n}$) throughout the experiment. Before the extraction setup, we predefined the alkaline extraction solution volume (5 L). We calculated the base solution (sodium hydroxide, 5 M), supplemented it, and added it to the known volume of the alkaline extraction solution of the prior sampling date. We exchanged the alkaline extraction solution when the volume exceeded the vessel capacity. After exchanging the extraction solution, we set the volume to the predefined value. We tested (data not shown) and found that the increase in volume was due to adding the alkaline solution to maintain the pH when the carboxylates were extracted.

$$V_{ex,n} = V_{ex,n-a} + \frac{W_{base,n} - W_{base,n-a}}{D_{base}} \quad \textbf{(Eq. S4)}$$

Where:

$V_{ex,n}$ , $V_{ex,n-a}$ = Volume of the alkaline extraction solution on the day n and n-a, L

$W_{base,n}$, $W_{base,n-a}$ = Weight of the base supplementation bottle on the day n and n-a, g

$D_{base}$ = Density of the base addition bottle, g $L^{-1}$

Volumetric production rate (mmol C $L^{-1}$ $d^{-1}$). The calculation of the volumetric production rate includes the extracted products and effluent product terms. As previously reported, we maintained the effluent product term (Palomo-Briones *et al.*, 2022). We modified the previously noted term of the extracted products to account for the change in volume and concomitant dilution of carboxylates throughout. We assumed that the system was at a steady state when taking samples. Therefore, we got:

$$Volumetric\ production\ rate = \frac{1}{V_{wet,x}}\left[\frac{C_{fe,n}V_{wet,x}}{HRT_n} + \frac{C_{ex,n}V_{ex,n} - C_{ex,n-a}V_{ex,n-a}}{t_n - t_{n-a}}\right] \quad \textbf{(Eq. S5)}$$

Where:

$C_{fe,n}$ = Concentration of the carboxylate in effluent on day n, mM C

$HRT$ = Hydraulic retention time on the day n, d

$C_{ex,n}$, $C_{ex,n-a}$ = Concentrations of the carboxylate in the extraction solution on days n and n-a, mM C

$V_{ex,n}$, $V_{ex,n-a}$ = Volume of the stripping solution on days n and n-a, L

$t_n$, $t_{n-a}$ = The day of operation n and n-a, d

Conversion efficiency: We calculated the conversion efficiencies to the product, for example, *n*-caproate, based on the total carbon feed. The conversion efficiency is independent of the changing wet working volume and therefore denotes the system performance with greater consistency throughout the experimental periods than the volumetric production rates.

$$EF_{product} = \frac{VP_{product}}{OLR} \times 100\% \quad \textbf{(Eq. S6)}$$

Where:

$VP_{product}$ = Volumetric production rate of product on the day n, mmol C $L^{-1}$ $d^{-1}$

$OLR$ = Total organic loading rate on the day n, mmol C $L^{-1}$ $d^{-1}$

**Total ammoniacal nitrogen**

The free ammonia (FA) concentration was determined based on total ammoniacal nitrogen, pH, and temperature using Eq. S7) (Zhen *et al.*, 2015).

$$Open \frac{[NH_3]}{[Total\ ammoniacal\ nitrogen]} = 1/\left(1 + \frac{10^{-pH}}{10^{-\left(0{:}09018+\frac{2729.92}{273.15+T}\right)}}\right) \quad \textbf{(Eq. S7)}$$

where $[NH_3]$ is the FA concentration (mg $L^{-1}$), [Total ammoniacal nitrogen] (mg $L^{-1}$) is the sum of the ammonium ion and free ammonia concentrations in the fermentation supernatant, and T is the temperature, °C

**Product-to-carboxylates specificity (% mol C)**

$$\frac{\gamma_p}{\sum_{i=1}^{n} \gamma_i} \times 100\% \qquad \textbf{(Eq. S8)}$$

$\gamma p$ = Production rate of a specific product, mmol C $L^{-1}d^{-1}$

$\gamma i$ = Production rate of all detected carboxylic acids, mmol C $L^{-1}d^{-1}$

### Results and discussion

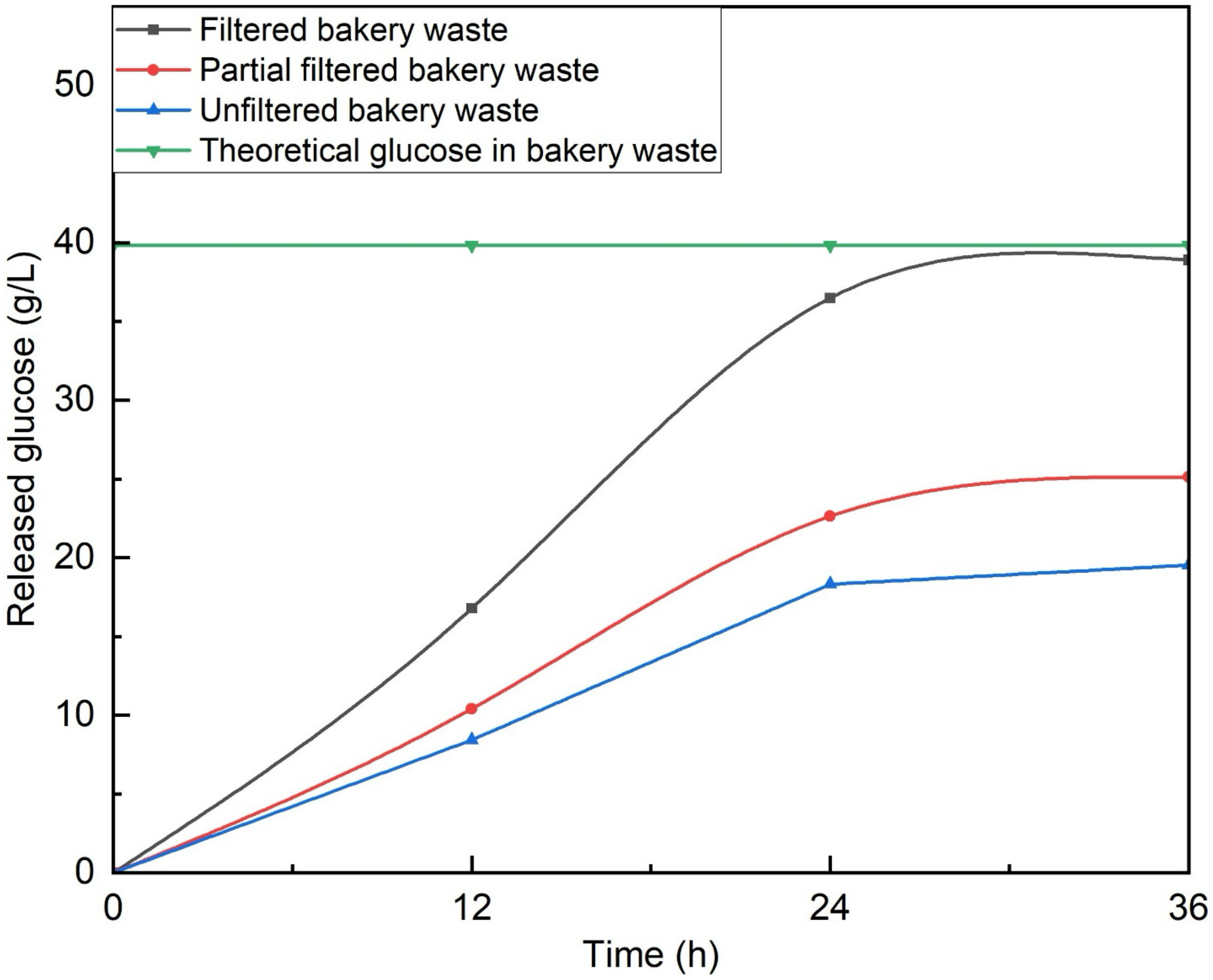


**Figure S3**. The time course profile of glucose released from the enzymatic hydrolysis of bakery waste.

**Table S3.** Optimization of bakery-waste mash to produce chain elongation precursors.

| | Parameters | | | Outcome | |
|---|---|---|---|---|---|
| ***Scenario*** | ***Mash with particles*** | ***Temperature ($^0$C)*** | ***Inoculum*** | ***Conc.Lactate (mM)*** | ***By-products*** |
| 1 | Large | 25 | Lactate microbiome | 103 +/- 32.8 | 3.00 % (Ethanol/Lactate) |
| 2 | Small | 25 | Lactate microbiome | 122 +/- 41.1 | 5.00 % (Ethanol/Lactate) |
| 3 | Medium | 25 | Lactate microbiome | 89.3 +/- 15.6 | 3.70 % (Ethanol/Lactate) |
| 4 | Large | 37 | Lactate microbiome | 98.7 +/-19.8 | 2.00 % (Ethanol/Lactate) |
| 5 | Small | 37 | Lactate microbiome | 159 +/- 48.2 | 4.00 % (Ethanol/Lactate) |
| 6 | Medium | 37 | Lactate microbiome | 104 +/- 12.6 | 3.50 % (Ethanol/Lactate) |
| 7 | Large | 50 | Lactate microbiome | 73.0 +/- 0.500 | 2.50 % (Ethanol/Lactate) |
| 8 | Small | 50 | Lactate microbiome | 84.4 +/- 0.510 | 2.70 % (Ethanol/Lactate) |
| 9 | Medium | 50 | Lactate microbiome | 76.3 +/- 14.1 | 3.00 % (Ethanol/Lactate) |
| 10 | Large | 25 | SiloSolve MC | 20.2 +/- 12.5 | Not measurable |
| 11 | Small | 25 | SiloSolve MC | 45.2 +/- 14.2 | Not measurable |
| 12 | Medium | 25 | SiloSolve MC | 37.3 +/- 8.90 | Not measurable |
| 13 | Large | 37 | SiloSolve MC | 30.4 +/- 10.6 | Not measurable |
| 14 | Small | 37 | SiloSolve MC | 14.7+/- 9.13 | Not measurable |
| 15 | Medium | 37 | SiloSolve MC | 36.4 +/- 13.5 | Not measurable |
| 16 | Large | 50 | SiloSolve MC | 11.8 +/- 0.0300 | Not measurable |
| 17 | Small | 50 | SiloSolve MC | 26.8 +/- 13.4 | Not measurable |
| 18 | Medium | 50 | SiloSolve MC | 8.92 +/- 2.46 | Not measurable |
| 19 | Large | 25 | *Bacillus coagulans* DSM1 | 36.3 +/- 11.6 | Not measurable |
| 20 | Small | 25 | *Bacillus coagulans* DSM1 | 55.2 +/- 10.2 | Not measurable |
| 21 | Medium | 25 | *Bacillus coagulans* DSM1 | 49.6 +/- 0.460 | Not measurable |
| 22 | Large | 37 | *Bacillus coagulans* DSM1 | 58.6 +/- 12.6 | Not measurable |
| 23 | Small | 37 | *Bacillus coagulans* DSM1 | 102 +/- 23.4 | Not measurable |
| 24 | Medium | 37 | *Bacillus coagulans* DSM1 | 98.1+/- 14.4 | Not measurable |
| 25 | Large | 50 | *Bacillus coagulans* DSM1 | 136 +/- 15.4 | Not measurable |
| 26 | Small | 50 | *Bacillus coagulans* DSM1 | 263 +/- 0.120 | Not measurable |
| 27 | Medium | 50 | *Bacillus coagulans* DSM1 | 169 +/- 0.240 | Not measurable |
| | | | | ***Conc. Ethanol (mM)*** | |
| 28 | Large | 30 | Kornbrand ``PREMIUM`` | 38.5 +/- 1.86 | $CO_2$ and Trace acetate |
| 29 | Small | 30 | Kornbrand ``PREMIUM`` | 180 +/- 0.120 | $CO_2$ |
| 30 | Medium | 30 | Kornbrand ``PREMIUM`` | 112 +/- 1.27 | $CO_2$ and Trace acetate |

### Result summary of the ethanol-bioreactor

Contrary to the lactate-bioreactor, we initiated the production of *n*-caproate in the ethanol-bioreactor by adding acetate as an electron-acceptor. The result of the ethanol-bioreactor showed that throughout the whole chain elongation period (Period-1 and Period-2), a mixture of the short-chain carboxylates and methane production dominated other products, a typical example of the anaerobic digestion system (Nguyen *et al.*, 2019) (**Fig. S4**).

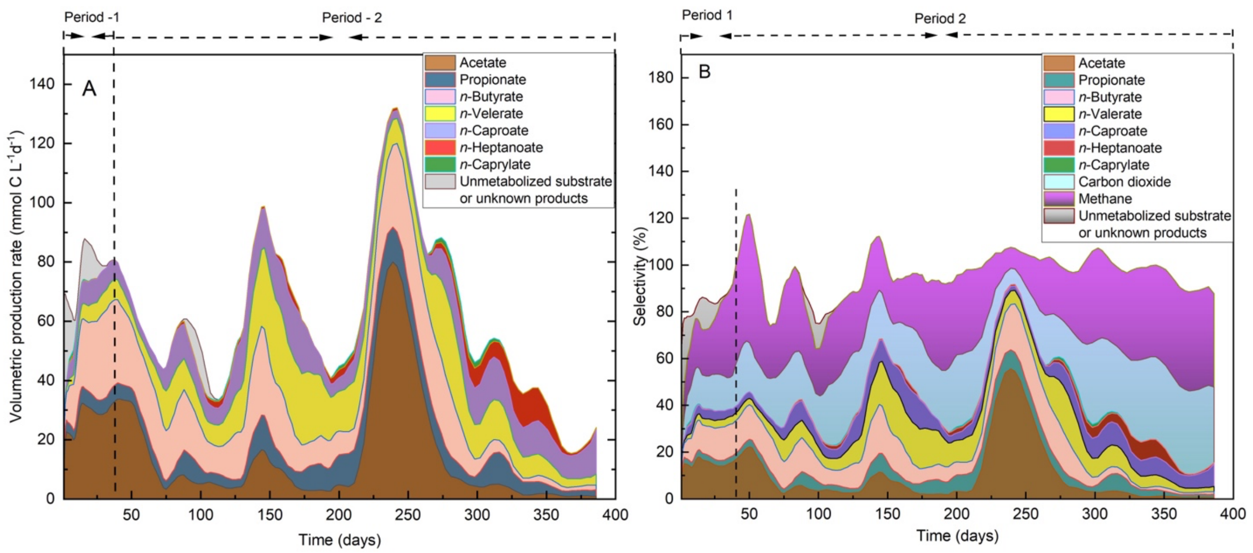


**Figure S4**. Carboxylate, unmetabolized substrate/unknown product, and gas profiles throughout the operating period from the bioreactor that was fed with ethanol as an intermediate after pre-treatment of bakery waste: **A)** volumetric production rates, and **B)** selectivity. The volumetric production rates and selectivities were calculated as a six-measurement moving average. The operating Periods I (a) – II (b) are indicated and described in the Materials and Methods section.

### Sankey diagram for the ethanol-bioreactor chain elongation system

The Sankey diagram with the carbon flow in our bakery-waste fermentation system provides a summary and clear visual representation of microbial chain elongation with (**Fig. S5A**) and without (**Fig. S5B**) acetate as the external electron acceptor. We normalized all bakery waste TOC measured values to 100 kg of carbon atoms (kg C) or 143 kg of dried bakery waste. The bakery waste microbial chain-elongation with acetate as an externally added electron acceptor resulted in 2.77 kg C of *n*-caproate, 0.198 kg C of *n*-caprylate, 0.152 kg C of *n*-heptanoate, 26.7 kg C of SCCs, 62.3 kg C of gas released to the atmosphere, which predominantly consists of carbon dioxide, and 36.7 kg C of unmetabolized substrate or unknown products (**Fig. S5A**). However, the microbial chain elongation without acetate addition resulted in 3.46 kg C of *n*-caproate, 0.114 kg C of *n*-caprylate, 0.836 kg C of *n*-heptanoate, 17.9 kg C of SCCs, 61.8 kg C of gas released to the atmosphere (predominantly carbon dioxide), and 15.45 kg C of unmetabolized substrate or unknown products (**Fig. S5B**).

The summary results of ethanol-rich substrate from bakery waste chain elongation technology suggested that the production of *n*-caproate and *n*-caprylate is not feasible. We could not change performance even during a lengthy operating period, including several operating optimizations. Possibly, the ratio of ethanol to other substrates was too low to produce MCCs, lacking enough reducing power.

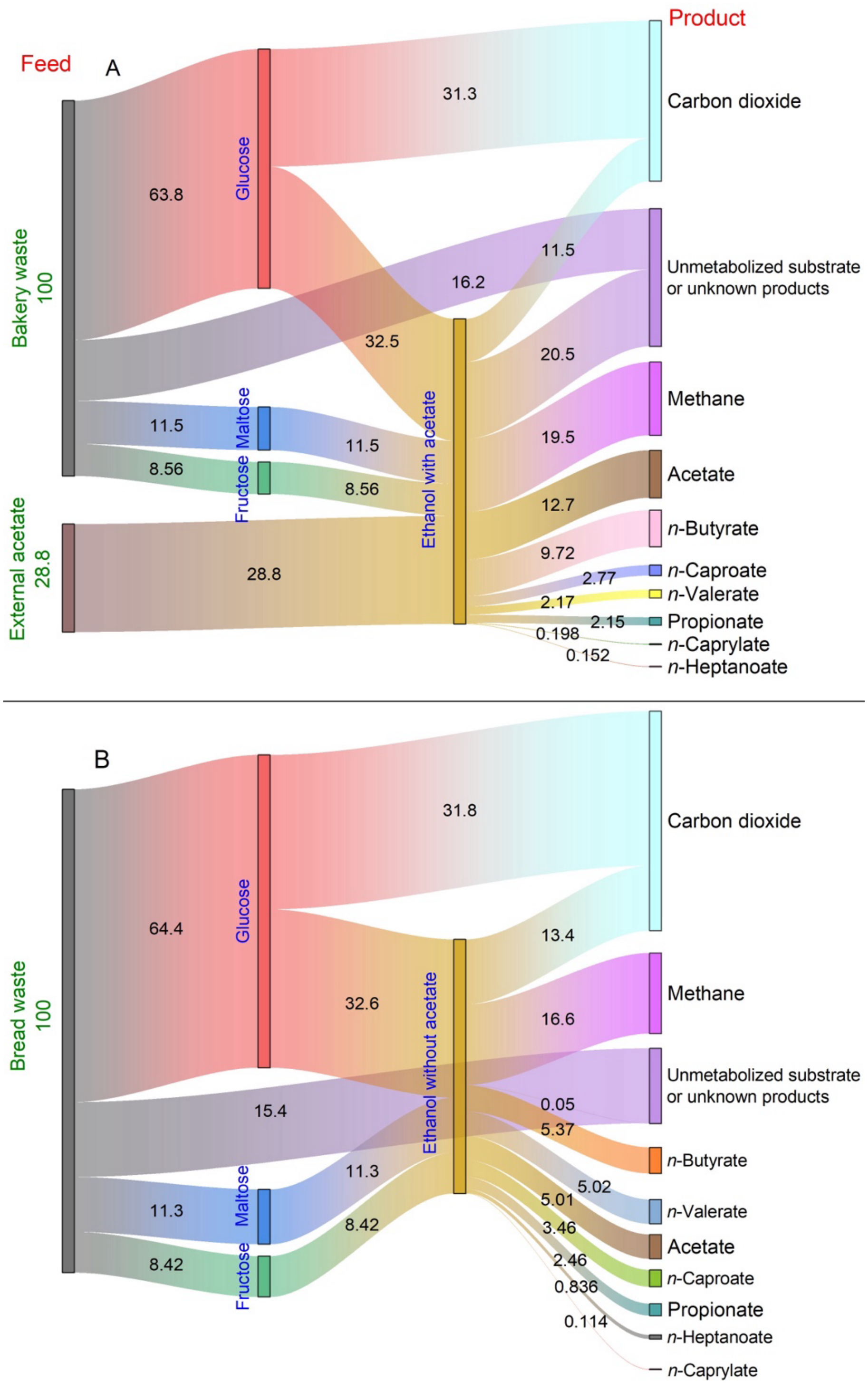


**Figure S5**. Sankey diagram of the carbon cradle-to-gate flows in the ethanol-bioreactor chain elongation system: A) chain elongation with the supply of external acetate, and **B)** chain elongation without the supply of external acetate.

**The choice of inoculum**

We utilized sludge from a manure-treating anaerobic digester and sludge from an anaerobic digestion at a municipal wastewater treatment plant to choose the best inoculum for the bakery waste chain-elongation process. We focused on producing *n*-caproate and hydrogen gas during the test and monitored them daily. First, *n*-caproate was the desired end-product of the bakery waste project and second, hydrogen gas might indicate the stability of the chain-elongation process. The low partial pressure of $H_2$ leads to the oxidation of SCCs and MCCs through the $\beta$-oxidation pathway (Spirito *et al.*, 2014). Furthermore, when the $H_2$ partial pressure is lower than 60 $Nm^{-2}$, the reducing power of NADH is used for the reduction of $H^+$ to produce $H_2$, rather than binding acetyl-CoA to the existing acyl-CoA chain (*i.e.*, acetyl-CoA or butyryl-CoA) (Angenent *et al.*, 2004). Therefore, we measured and monitored *n*-caproate and hydrogen. After one week, we reached a concentration of 34.3, 38.8, and 51.3 mM of *n*-caproate for sludge from a manure-treating anaerobic digester, anaerobic digestion sludge from a municipal wastewater treatment, and a mixed sludge of the two inocula, respectively. However, hydrogen amounts were similar during the first 4 days of the inoculum test in the continuous chain-elongation process for all inocula, and the bioreactor with inoculum from manure biogas produced approximately 48% less hydrogen than the others. Therefore, we chose a combination of manure biomass and municipal wastewater treatment inocula for the subsequent chain-elongation process.

### Pseudo-granules

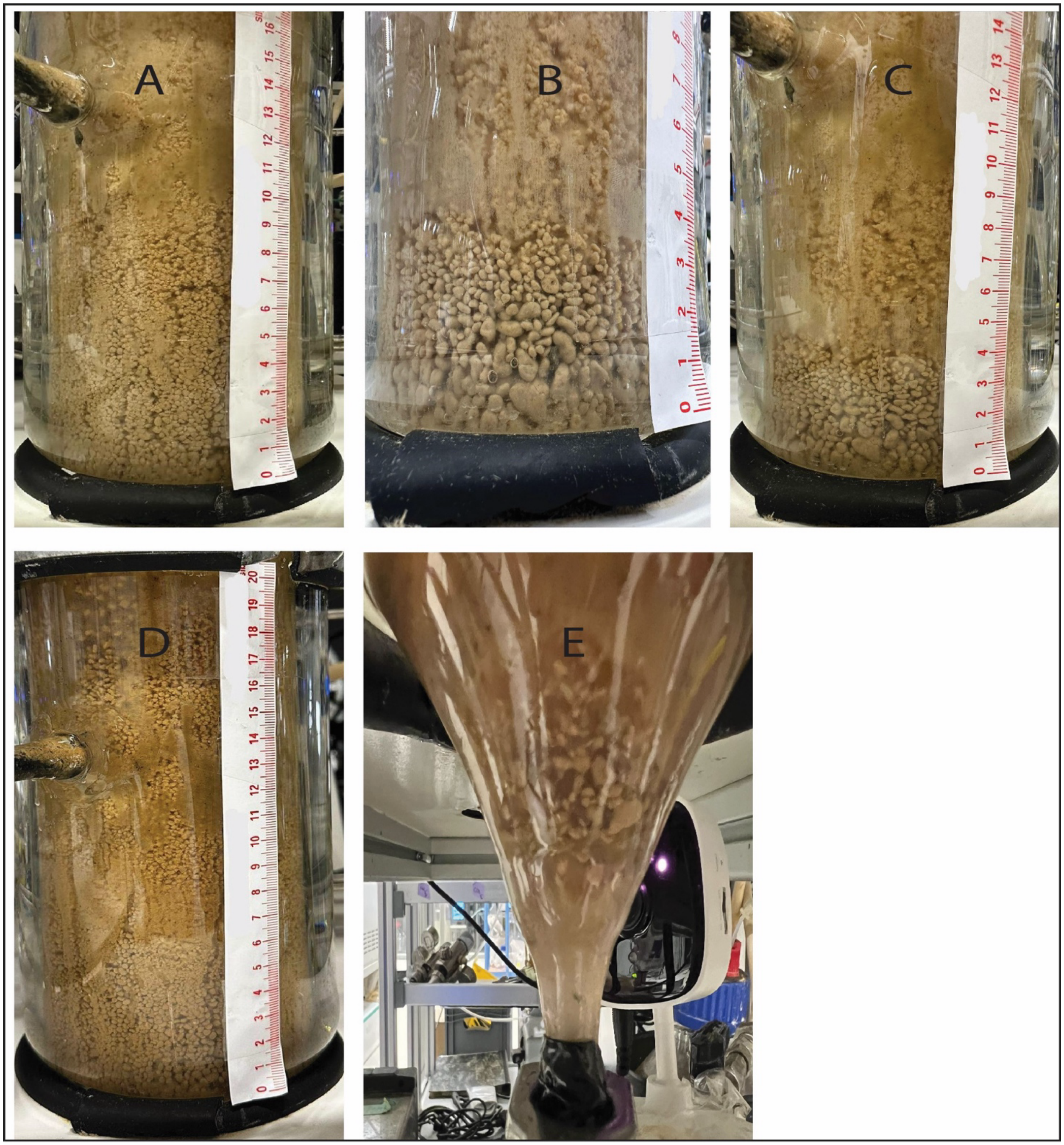


**Figure S6.** Pseudo-granules in the bakery waste chain-elongation bioreactor. The granules appeared and disappeared. An example of the pseudo-granule disappearance process (Pictures A, B, C, D, and E).
